# Vergence accuracy in an autostereoscopic display

**DOI:** 10.1101/2021.07.29.454355

**Authors:** Luca Lo Verde, Anthony M. Norcia

## Abstract

When fixating an object, observers typically under or over-converge by a small amount, a phenomenon known as “fixation disparity”. Fixation disparity is typically measured with physical fixation targets and dichotically presented nonius lines. Here we made fixation disparity measurements with an autostereoscopic display, varying the retinal eccentricity and disparity of the fixation targets. Measurements were made in a group of four practiced observers and in a group of thirteen experimentally naïve observers. Fixation disparities with a zero-disparity target were in the direction of fixation behind the plane of the screen and the magnitude of the fixation disparity grew with the eccentricity of the fixation targets (1-5 deg in the practiced observers and 1 – 10 deg in the naïve observers). Fixation disparity also increased with increasing disparity of the targets, especially when they were presented at crossed disparities. Fixation disparities were larger overall for naïve observers who additionally did not converge in front of the screen when vergence demand was created by crossed disparity fusion locks presented at 5 and 10 deg eccentricities.

## Introduction

Observers with normal binocular vision typically fixate slightly in front of or behind an objective of regard, a phenomenon known as fixation disparity (Hoffmann and Bielschowsky, 1900; Ogle et al., 1967). Fixation disparity is typically measured using dichoptic nonius lines to read-out the perceived visual direction of each eye and this measurement is generally well correlated with the pointing direction of the two eyes in observers with normal binocular vision (Hillis and Banks, 2001).

Fixation disparity can be exaggerated by imposing different degrees of retinal disparity on display elements. In the classical studies of fixation disparity, these image disparities were produced through the use of prisms (Ogle et al., 1967). Fixation disparity has also been reported to increase with increasing retinal eccentricity of disparate targets (Carter, 1964; Debysingh et al., 1986; Ukwade, 2000).

Measures of fixation disparity are commonly measured clinically with physical cards or devices and prisms (Mallett, 1964; Sheedy, 1980; Wesson and Koenig, 1983). Here we studied fixation disparity using a large-format lenticular autostereoscopic display in practiced and experimentally naïve observers. Changes on vergence demand were induced by disparate targets presented over a range of retinal eccentricities. Fixation disparity increased with both increasing vergence demand and retinal eccentricity of the stimulus to fusion, especially for crossed disparities (increased convergence demand) and for experimentally naïve observer

## Methods

### Participants

Two groups of observers participated. Four highly practiced participants (2 females, mean age: 42 ± 18 years; range 31-69; two observers were co-authors of this work) were tested to tune the experimental parameters. These observers had normal- or corrected-to-normal vision, and normal stereoacuity.

A second, experimentally naive group of thirteen participants (7 females, mean age: 19.9 ± 0.8 years; range: 19-21 years old) were tested after being screened for having normal or corrected to normal visual acuity and stereoacuity (Bailey Lovie Eye Chart; Randot Stereotest). Consenting and experimental procedures were approved by the Institutional Review Board of Stanford University. The research conformed to the tenets of the Helsinki Convention.

### Apparatus and Stimuli

The experiment was performed in a quiet room in total darkness. The stimuli were developed in MATLAB (The Mathworks, Inc., Natick, MA) using Psychtoolbox-3 (Brainard, 1997; Pelli and Vision, 1997; Kleiner et al., 2007) and delivered using a SeeFront autostereoscopic 3D monitor (SeeFront GmbH, Frankfurt, Germany; model SF3D320-MP). A schematic representation of the stimulus configuration, depicted as a red/blue anaglyph for visualization purposes, is shown in Figure 1A. Nonius lines were used as a high-precision subjective indicator of vergence posture (McKee and Levi, 1987). The dichoptic nonius lines were gray, 21 minutes of arc long and were vertically separated by 36.5 minutes of arc. The top nonius lines was delivered to the right eye and fixed in position, always presented in the center of the screen and in the center of the right eye fusion lock component. The participant could adjust the horizontal position of the other nonius line using a track wheel device. The initial position of the adjustable nonius line varied pseudo-randomly on each trial, with a maximum possible displacement of 14.4 arcmin from the screen center. The background was black.

**Figure 1.**
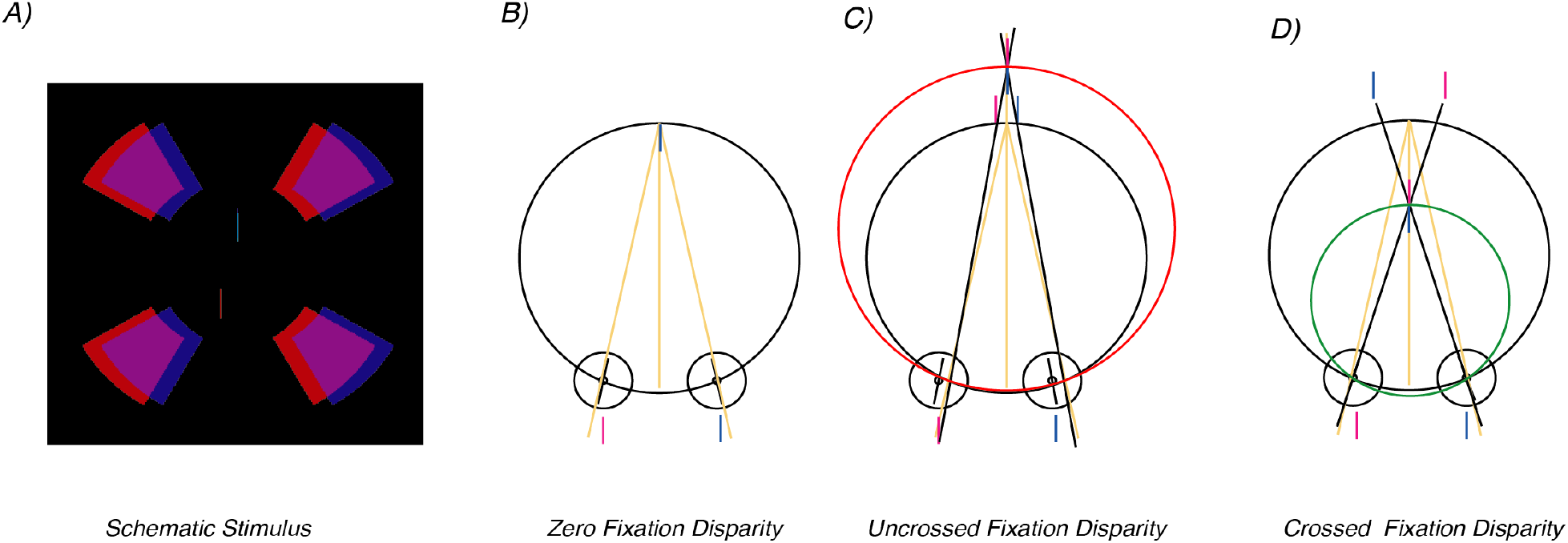
A) Schematic representation of the stimuli, in this case depicting the dichoptic fusion lock with 1 deg of eccentricity and 0.2 degrees of uncrossed disparity. In this anaglyphic representation, the magenta portions of the stimuli are delivered to the left eye while the blue ones are seen by the fellow eye (every component of the stimulus was gray in the actual experiment). We manipulated both eccentricity and disparity of the fusion lock surrounding the nonius lines. The upper nonius line was always a fixed reference, while the participants could adjust the horizontal position of the lower, probe nonius line to perceptually align them. B) Zero fixation disparity. Physically aligned nonius lines at the plane of the screen (black curve) are subjectively aligned with zero nonius offset. C) Uncrossed fixation disparity. The eyes are diverged relative to the plane of the screen, subjectively aligned targets have a negative nonius offset. D) Crossed fixation disparity. The eyes are converted relative to the plane of the screen, subjectively aligned targets have a positive nonius offset.

The binocular fusion lock consisted of a radial pattern composed of four circular sectors surrounding the nonius lines. The fusion lock could be placed at different retinal eccentricities around the nonius stimuli: four eccentricities for the pilot data collection (1°, 1.25°, 2.5° and 5° of eccentricity) and three eccentricities for the main data pool (1°, 5° and 10° of eccentricity). These eccentricities were defined as the distance from the screen center and the inner radius of the four circular sectors around the nonius lines. The circular sector sizes were scaled to equate their visibility according the cortical magnification factor (Baseler et al., 1994) and thus the fusion locks presented at higher eccentricities were larger than the ones presented closer to the center of the visual field. The equation used to estimate the cortical area *A* subtended by circular sectors is the following:

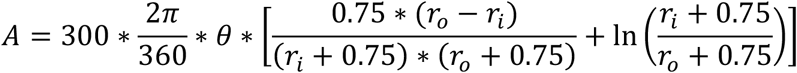

Where *θ* represents the angle of a given circular sector, *r*_*i*_ represents the inner radius between the screen center and the inner margin of each circular sector, and *r*_*o*_ represents the outer radius between the screen center and the outer margin of each circular sector. The fusion lock was presented at thirteen different horizontal disparities (6 crossed and 6 uncrossed disparities plus zero disparity: 0°, ±0.05°, ±0.1°, ±0.2°, ±0.4°, ±0.6° and ±0.8°).

### Procedure

Participants sat comfortably 70 cm from the monitor. They held a rotary USB input device used for collecting responses. Experimentally naive participants were familiarized with the stimuli and task by having them perform 5 training trials. Practiced observers had extensive experience with the task prior to data collection. Each trial began with a blank, dark screen, followed by the appearance of the fusion lock stimulus, presented at zero disparity for 750 msec. Subsequently, the fusion lock assumed the designated disparity for that trial and the nonius lines appeared in the center of the screen. The participants were asked to focus on the central nonius lines and to ignore the surrounding radial fusion lock pattern. These stimuli remained visible on the screen until the end of the trial.

The participants used the rotary input device to adjust the position of the probe nonius line until they perceived the two nonius lines as vertically aligned. The device allowed them to progressively and smoothly move the probe nonius line in either horizontal direction (step size: 1 pixel ≈ 1.80 minutes of arc) to align the vertical nonius lines. The participants pressed a dedicated button on the response device to proceed to the next trial once they were satisfied with the perceived alignment of the nonius lines.

Figure 1B illustrates the case of zero fixation disparity. Here physically aligned nonius lines are seen as subjectively aligned, indicating that vergence was on the plane of the screen, indicated by the black Vieth-Müller circle presented as an approximation of the empirical horopter. Fig. 1C indicates the case of uncrossed fixation disparity where the eyes are diverged relative to the plane of the screen and subjective alignment occurs with a negative nonius offset, under our convention. Fig. 1D indicates the case of a crossed fixation disparity where the eyes are converged relative to the plane of the screen and subjective alignment occurs with a positive nonius offset, under our convention.

There were no time constraints on the perceptual alignment task, and the participants were instructed to achieve a stable alignment before finishing a given trial. Each experimental session consisted of 6 trial repetitions for each of the 3 fusion lock eccentricities (4 fusion lock eccentricities for the experienced observers pool) and the 13 fusion lock disparities, for a total of 246 trials per participant, split in 3 separate blocks each consisting of 82 trials.

### Conventions

We adopted the disparity conventions in widespread use across classical optometry works (Ogle et al., 1967) by assigning uncrossed fusion lock disparities to the positive range of the presented disparity axis while the crossed disparity fusion locks are assigned negative values. We first computed deviations from each presented vergence demand by subtracting the actual vergence demand from the final adjusted position of the participants-controlled nonius line which had to be perceptually aligned with the other, fixed nonius line. Therefore, a positive (negative) deviation indicates a failure to converge (diverge) the eyes enough to have each dichoptic fusion mask lie on each corresponding points on the two retinae. We computed these deviations for the three fusion lock eccentricities as a function of vergence load, as well as having the deviations for each vergence load as a function of fusion lock eccentricity to better visualize the influence of these variables on the task.

## Results

The relative alignment of the nonius lines was continuously tracked until they were perceptually aligned. Figure 2 A-D shows example time-courses of the alignment judgements for each of the fusion lock eccentricities for the zero-disparity fusion lock condition (1.0, 1.25, 2.5 and 5 deg for the practiced observers pool and 1.0, 5 and 10 deg for the naïve participants pool, respectively). The data are from the four practiced observers in panels A-D and the thirteen naïve observers in E-G. The zero-disparity condition represents a conventional estimate of fixation disparity, but with eccentric fixation locks. The average fixation disparity increased with the eccentricity of the fusion lock (dashed line), as did the variability of the setting (red band). The dashed red lines indicate the average of the final adjustment before trial end for each participant, and the red shade represents the standard error of the means.

**Figure 2.**
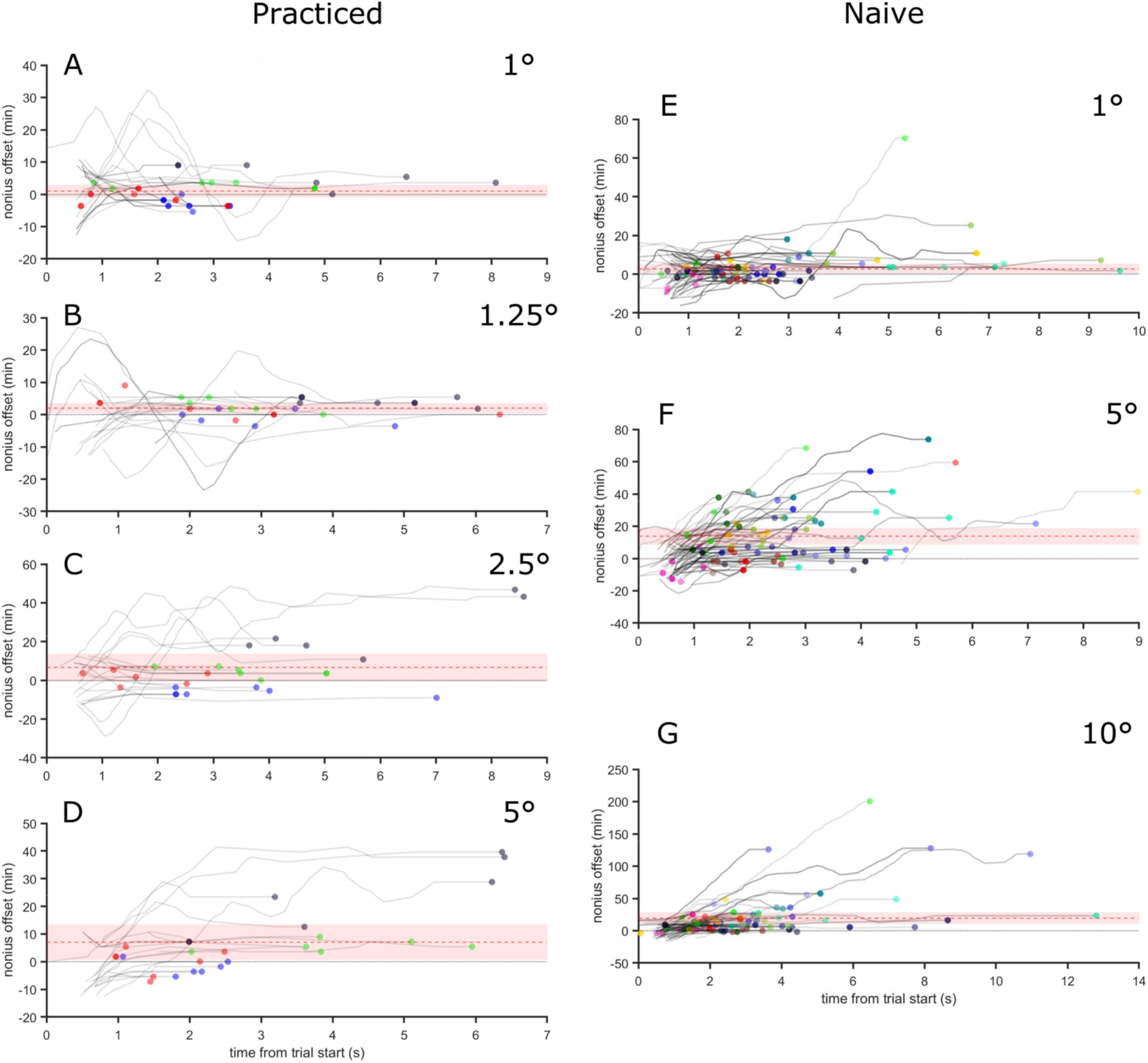
Time course of the nonius position adjustments from trial onset, plotted for each fusion lock eccentricities (A: 1°; B: 1.25°; C: 2.5°; D: 5°) for the zero-disparity condition for the practiced participant pool. Each color represents a single participant, the red dashed line represents the average of the final nonius position across all participants, the light red area defines the standard error of the mean. E-G) data from the 13 naïve participants using the same convention.

The full data sets over all fixation lock disparities and eccentricities are shown in Figure 3. The dashed diagonal lines indicate the imposed disparity of the fusion lock and thus the vergence demand. For both practiced and naïve observers, the nonius offset for subjective alignment accurately tracks the disparity of the low-eccentricity, uncrossed fusion locks (1 deg in both groups (light blue curves in both A and B) and additionally at 1.25 deg eccentricity in the practiced group but undershoots it for crossed disparities especially in the naïve observers (compare Fig. 3 A to B). As the eccentricity of the fusion lock increases to 2.5, 5 and 10 deg eccentricity, the nonius offset setting increasingly fails to track the disparity of the fusion lock, especially for crossed disparities (see Fig. 2 C, D and F, G). The divergence of the nonius settings from the imposed demand implies that the eyes are less converged or diverged than the demand imposed by the disparity of the fusion lock. Notably, in the naïve observers, the nonius offset for crossed fusion locks are on the uncrossed side for the largest fusion lock disparities.

**Figure 3.**
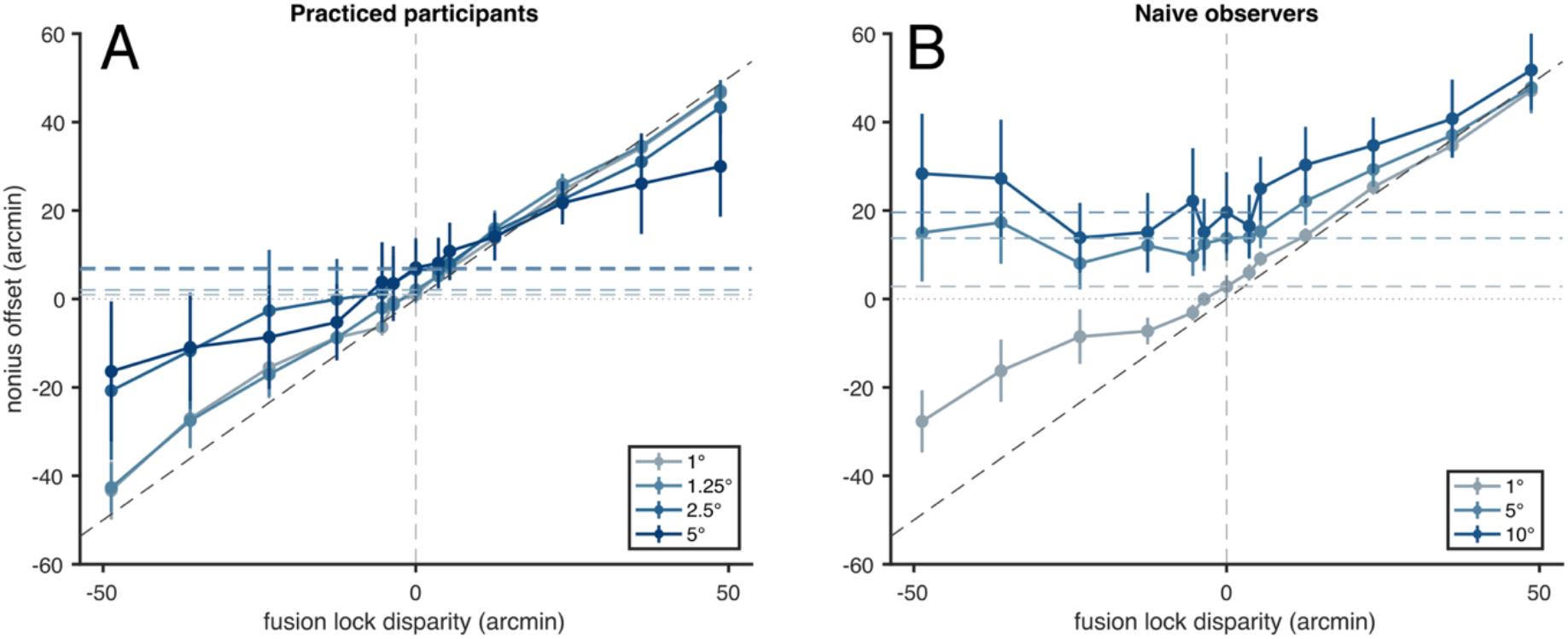
Nonius offset settings for practiced (A) and Naïve observers (B). Eccentricities in degrees are reported in the legends.

To better visualize the magnitude of the biases, we transformed the nonius offsets in deviations from imposed vergence load and visualized the data of Figure 3 in two ways: first as a function of fusion lock disparity, with eccentricity as the parameter and then as a function of disparity with eccentricity as the parameter. At zero disparity and 1 degree of eccentricity, the fixation disparity in the practiced observers was 0.98 ± 1.97 arcmin and in the naïve observers it was 2.82 ± 2.6 arcmin. At 5 deg, where both groups provided data, these values were 7.06 ± 6.44 arcmin and 13.73 ± 5.1 arcmin, respectively. At 10 degrees of eccentricity (for the naïve participants pool), the fixation disparity was 19.58 ± 9.13 arcmin.

As the disparity of the fusion lock increases, nonius offset deviation remains essentially constant for all uncrossed disparities when the fusion lock eccentricity is 1 deg in both groups and 1.25 deg in the practiced observers. At 2.5 deg and higher fusion lock eccentricities, the zone where fixation disparity is small shrinks to disparities less than ≈5.41 arcmin. The fixation disparity increases with eccentricity for all disparities, especially for crossed disparities and larger disparity magnitudes in both groups. The asymmetry in settings between crossed and uncrossed disparities is readily apparent when deviation is plotted as a function of eccentricity for the two disparity signs (Fig 4 C, D).

**Figure 4.**
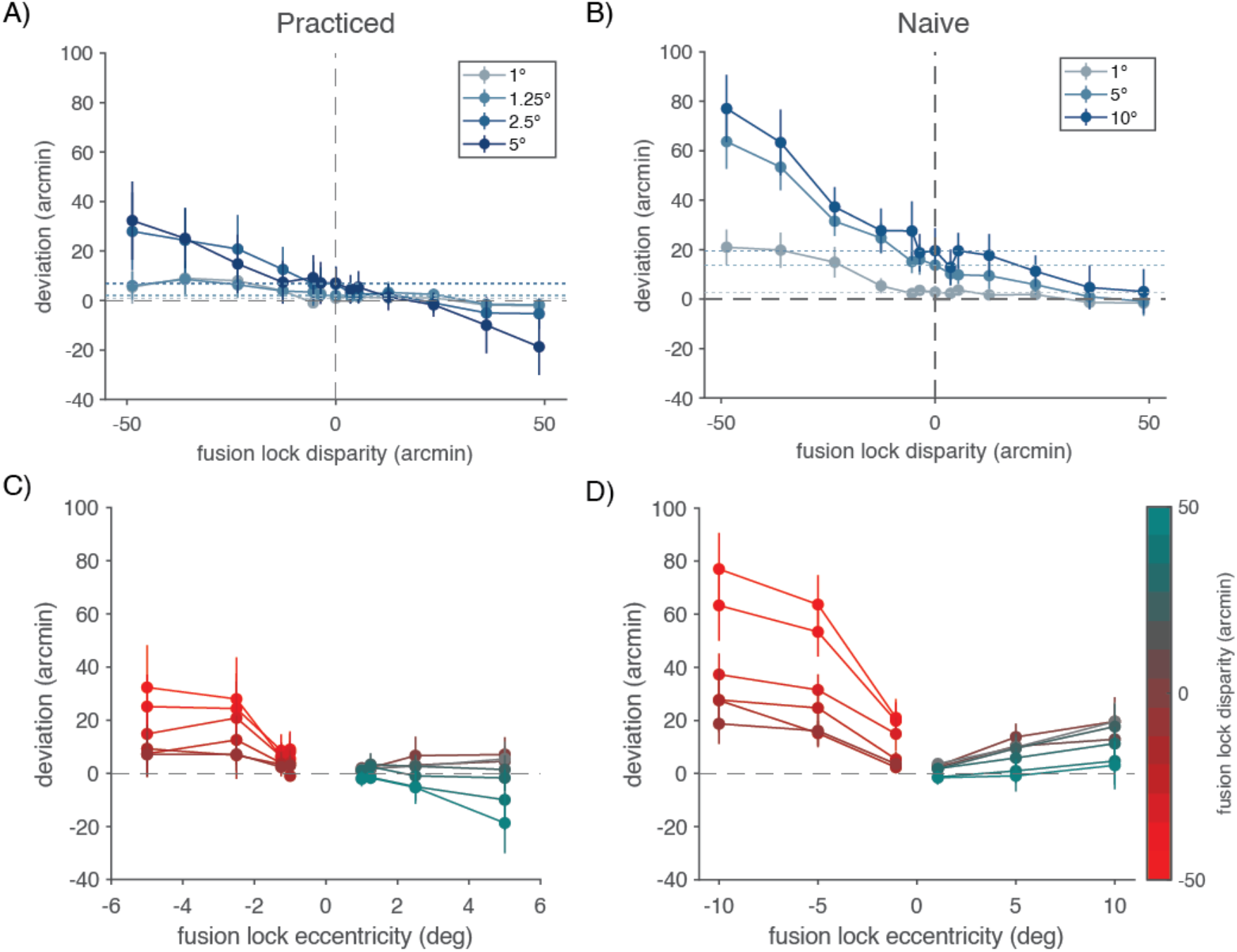
Deviations of settings from vergence demand. A) Deviation as a function of fusion lock disparity for practiced observers. Deviation grows with increasing fusion lock, especially for larger crossed disparities. B) Deviation as a function of fusion lock disparity for naive observers. Trends in experience observers are exaggerated. C) Deviation as a function of fusion lock eccentricity for practiced observers. Deviation increases for more eccentric fusion locks, especially for crossed disparity. D) Deviation as a function of fusion lock eccentricity for naïve observers. Trends in practiced observers are exaggerated in the naïve observers. For visibility purposes, we translated the negative fusion lock disparities curves (plotted in shades of red) to negative eccentricities in C and D.

## Discussion

Our results with an autostereoscopic display qualitatively recapitulate results from the clinical and psychophysical literatures. We find that the degree of nonius misalignment increases with increasing disparity of the fusion locks. Increasing the disparity of the fusion lock mimics the addition of prisms as used in clinical studies or the use of mirror deflectors (Schor et al., 1986), but does not induce a shift of all visible elements over the visual field. In our case, only the fusion lock stimuli varied in disparity, but the monitor bezel and the dimly lit room surround did not.

In both observer groups, fixation disparity measured with a zero-disparity fusion lock is positive, meaning that the eyes are converged behind the plane of the display. The magnitude of the fixation disparity increases in both groups as the fusion lock eccentricity increases, as observed in previous studies (Carter, 1964; Debysingh et al., 1986; Ukwade, 2000).

As increasing disparity is introduced into the fusion lock, the nonius offset directly tracks the magnitude of the imposed disparity for uncrossed fusion locks when the eccentricity of the fusion lock is low (*e.g*. 1 to 1.25 deg). The uncrossed fusion locks are portrayed behind the plane of the monitor. At larger eccentricities, the nonius offset fails to track the imposed disparity by increasing amounts in the practiced observers (see Fig 3C and D). In the naïve observers the nonius offset matches the largest of the imposed uncrossed disparities, but over-compensates in the uncrossed direction for smaller fusion lock disparities (see Fig. 3 F and G).

The crossed disparity fusion locks render the images in front of the plane of the monitor. In both observer groups, for the smallest fusion lock eccentricities, the nonius offset tracks the sign of the disparity, but fails to keep up with the magnitude of the disparity (see Fig. 3 A,B and E). In the practiced group, the function is shallower than the imposed disparity demand for both signs of disparity and by 5 deg of eccentricity, the effect is approximately symmetric for crossed and uncrossed disparities. In the naïve observers, the presentation of crossed disparity fusion locks fails to drive convergence in front of the plane of the monitor at 5 and 10 deg eccentricities of the fusion lock – fusion lock disparity has no effect and does not alter the fixation disparity value measured for zero disparity.

Previous research on the perception of depth from disparity has compared stereoacuity for simple real-world targets and simulated versions of them presented in a stereoscope (McKee and Taylor, 2010). For two unpracticed observers, depth thresholds in the stereoscope were much worse than their real-space depth thresholds. Superiority of cue integration has also been reported for real versus stereoscopically simulated displays (Buckley and Frisby, 1993). The autostereoscopic and other types of dichoptic viewing system inherently present conflicts between disparity cues and accommodation/proximity cues and this may affect the perception of depth from the fusion locks. However, this can’t be the whole explanation as cue conflict is equal over the different fusion lock eccentricities. In the naïve observers, performance is relatively good for the 1 deg fusion lock eccentricity, but not for the 5 and 10 deg eccentricities. This suggests another factor – peripheral disparity sensitivity – may be at play. Peripheral stereoacuity can improve more than central visual acuity with practice, at least in some observers (Fendick and Westheimer, 1983) and it is possible that the naïve observers are not as able to use peripheral disparity information in the context of our experimental conditions.

## Acknowledgments

Funded by a Stanford University Bio-X Interdisciplinary Initiatives Seed Grants Program (IIP) grant.

## Notes

### Competing Interest Statement

The authors have declared no competing interest.

